# Pleiotrophin signals through ALK receptor to enhance growth of neurons in the presence of inhibitory CSPGs

**DOI:** 10.1101/2022.04.01.486745

**Authors:** Somnath J Gupta, Matthew A Churchward, Kathryn G Todd, Ian R Winship

**Affiliations:** Neurochemical Research Unit, Department of Psychiatry, Faculty of Medicine and Dentistry, University of Alberta, 12-127 Clinical Sciences Building, Edmonton, AB T6G 2R3, Canada; Neuroscience and Mental Health Institute, University of Alberta, Edmonton, AB T6G 2R3, Canada; Department of Biology and Environmental Sciences, Concordia University of Edmonton, T5B4E4, Edmonton, AB, Canada

**Keywords:** Plasticity, neurons, CSPGs, PTN, ALK, neuronal growth

## Abstract

Chondroitin sulfate proteoglycans (CSPGs), one of the major extracellular matrix components of the glial scar that surrounds central nervous system (CNS) injuries, are known to inhibit the regeneration of neurons. This study investigated whether pleiotrophin (PTN), a growth factor upregulated during early CNS development, can overcome the inhibition mediated by CSPGs and promote the neurite outgrowth of neurons *in vitro*. The data showed that a CSPG matrix inhibited the outgrowth of neurites in primary cortical neuron cultures compared to a control matrix. PTN elicited a dose dependent increase in the neurite outgrowth even in the presence of the growth inhibitory CSPG matrix, with optimal growth at 15 ng mL^-1^ of PTN (114.8% of neuronal outgrowth relative to laminin control). The growth promoting effect of PTN was blocked by inhibition of the receptor anaplastic lymphoma kinase (ALK) by alectinib in a dose dependent manner. Neurite outgrowth in the presence of this CSPG matrix was induced by activation of the protein kinase B (AKT) pathway, a key downstream mediator of ALK activation. This study identified PTN as a dose-dependent regulator of neurite outgrowth in primary cortical neurons cultured in the presence of a CSPG matrix, and identified ALK activation as a key driver of PTN-induced growth.

**Summary statement:** Function in the central nervous system (CNS) is attributed to the complex interactions of neurons and glia. These cells are anchored in extracellular matrix (ECM) which constitutes about 10% - 20% of brain volume. Cells in the brain produce different components of the ECM in brain including chondroitin sulfate proteoglycans (CSPGs). After a nervous system injury, glial cells produce excess CSPGs that restrict the regeneration of neurons, thus limiting functional recovery. This study examines the role of the endogenous growth factor pleiotrophin (PTN) in driving the growth of neurons even in the presence of inhibitory CSPGs, and anaplastic lymphoma kinase (ALK) receptor as a key mediator by which PTN potentiates growth.

## Introduction

Chondroitin sulfate proteoglycans (CSPGs) are a major component of the extracellular matrix that surrounds cells of the central nervous system (CNS) (Siebert, Conta Steencken and Osterhout, 2014). CSPGs maintain CNS health by regulating growth of axons during development and protecting against oxidative stress (Morawski *et al*., 2004). After CNS injury, activated glial cells increase synthesis of CSPGs to form a glial scar that contains the injury site (Siebert, Conta Steencken and Osterhout, 2014). The scar reduces lesion growth but also acts as a barrier for the regeneration of neurons that may limit functional recovery (Siebert, Conta Steencken and Osterhout, 2014; Wang *et al*., 2009). CSPGs are also upregulated in neurodegenerative conditions including Alzheimer’s disease (AD), Pick bodies in Pick’s disease, and Lewy bodies in Parkinson’s disease (PD) (Herradõn and Pérez-García, 2014). Several experimental approaches have been investigated to neutralize the growth inhibitory effect of CSPGs *in vivo*, including digestion of CSPGs by chondroitinase ABC (ChABC), knockdown of CSPG polymerisation enzymes by RNA interference (RNAi), and peptide blocking the signalling of two major receptors of CSPGs: protein tyrosine phosphatase sigma (PTPσ) and leukocyte antigen receptor (LAR) (Keough *et al*., 2016). Although these strategies have shown positive effects on the growth of neurons in model systems, there are disadvantages that may limit translation to clinical use. Both ChABC and siRNA are exogenous macromolecules that must be delivered directly to the injury site, have short half-lives *in vivo*, and could induce immunological response due to repeated administration (Wong, 2008). Potentiating regeneration of neurons without degrading the glial scar may be possible via the heparin-binding growth factor pleiotrophin (PTN). PTN has long half life (Dreyfus *et al*., 1998; Paveliev *et al*., 2016) and may overcome the disadvantage of repeated administration associated with the use of ChABC and siRNA. PTN is a developmentally regulated protein whose expression peaks (in rats) 3-4 weeks after birth (Wang, 2020). Notably, PTN expression is low in adults but increases transiently after CNS injury (Wang, 2020). PTN is associated with neuroplasticity including the maturation of new neurons and induction of neurite outgrowth (Tang *et al*., 2019). Notably, PTN may reverse aggrecan (a prominent CSPG) mediated inhibition *in vitro* by preventing the binding of (Paveliev *et al*., 2016). In PD, PTN has shown to reduce nigrostriatal degeneration and improve functional recovery (Herradõn and Pérez-García, 2014). Notably, PTN has a sustained presence in the tissue when injected into the nervous system directly. Moreover, identifying key receptors involved in PTN-induced neuroplasticity may allow for systemic pharmacotherapy.

PTN has several putative cell surface receptors, including receptor protein tyrosine phosphatase ζ (RPTPζ/PTPRZ), syndecans, nucleolin, neuropilin-1, integrin αVβ3 and αMβ2, N-syndecan receptor, glypican 2, neuroglycan-C, and anaplastic lymphoma kinase (ALK) (Paveliev *et al*., 2016; Wang, 2020). Moreover, PTN can integrate into the extracellular matrix, limiting CSPG interactions with growth-inhibitory receptor PTPσ for a sustained period (Paveliev *et al*., 2016).

This study investigates the potential role of ALK receptor activation by PTN to increase the growth of neurons in the presence of a CSPG matrix. ALK is expressed during early development in CNS, with low expression in the adult central nervous system (Vernersson *et al*., 2006; Iwahara *et al*., 1997). ALK signalling is critical for differentiation of neuronal progenitor cells to neurons and regulates their survival (Yao *et al*., 2013; Tang *et al*., 2019). By binding to CSPGs, PTN also reduces phosphatase activity of RPTPζ, which increases activity of ALK (Wang, 2020). Moreover, in developing neurons PTN signals directly via ALK receptor binding (Tang *et al*., 2019;Yanagisawa *et al*., 2009). Based on the evidence mentioned above ALK receptor plays a major role in neurogenesis and neuronal survival making it a potential receptor to investigate its involvement in PTN signaling in the presence of CSPGs. Thus, here we investigated to role of PTN signaling via ALK in driving neuron growth in the presence of inhibitory CSPGs. Our data reveals a dose dependent effect of PTN on neurite outgrowth in the presence of CSPGs. Notably, selective pharmacological inhibition of the ALK receptor attenuated the growth promoting effect of PTN.

## Results and Discussion

### Pleiotrophin induces neurite outgrowth in the presence of CSPGs

CSPGs are known to inhibit the growth of neurons. In order to understand the effect of PTN on neurite outgrowth, cortical neurons were cultured on the growth permissive (laminin) matrix or on the growth inhibitory (laminin + CSPGs) matrix and treated with different concentrations of PTN (5 ng mL^-1^ – 20 ng mL^-1^) for 72 h. MAP2 immunofluorescence was used to quantify neurite outgrowth. Neurons showed extensive neurite outgrowth on the laminin matrix (Fig. 1A-C) that was inhibited by CSPGs (Fig. 1 D-F). PTN restored neurite extension in a dose dependent manner (Fig. 1G-I). Neuronal morphology was analysed using the Simple Neurite Tracker plugin for ImageJ (Longair, Baker and Armstrong, 2011) (Fig. 1J). A dose dependent effect of PTN on total neurite outgrowth was observed, with PTN counteracting the inhibitory effect of CSPGs at concentrations below 20 ng mL^-1^ and maximal effect at 15 ng mL^-1^ concentration (Fig. 1K). PTN treatment increased the complexity of cortical neuron neurite growth by increasing branched neurite growth, with maximum branch length observed at 15 ng mL^-1^ PTN (Fig. 1L), and increased the number of neurons with neurite outgrowth at 10-15 ng mL^-1^ PTN (Fig. 1M). This data suggests that PTN at specific concentrations potentiates neurite extension. PTN has been shown to promote cell survival signal in dopaminergic neurons invitro and in a mouse model of Parkinson’s disease PTN was shown to promote survival of grafted dopaminergic neurons, thus improving functional recovery of the nigrostriatal pathway (Herradõn and Pérez-García, 2014; Hida *et al*., 2007). These findings support PTN as a candidate to restore neurite extension in CSPG-rich lesion areas after CNS injury and other neurological conditions. Notably, PTN may also modulate the activity of glial cells including OPCs (Oligodendrocyte progenitor cell) and microglia. PTN induces differentiation of OPCs to mature oligodendrocytes, thus promoting myelination of developing neurons (Herradon and Ezquerra, 2009), and could therefore potentiate remyelination after injury. Microglia increase their release of neurotrophic factors including ciliary neurotrophic factor (CNTF), nerve growth factor (NGF) and brain-derived neurotrophic factor (BDNF) after stimulation with PTN (Miao *et al*., 2012). Thus, in addition to direct actions on neurons, PTN is a strong candidate to generate an environment favouring the neuronal growth and its functionality by actions on multiple cell types. These data suggest PTN signaling may be an exciting approach to enhancing neuroplasticity after CNS injury, though the dose-response relationship for PTN in the presence of CSPGs will be important to verify *in vivo*.

**Figure 1:**
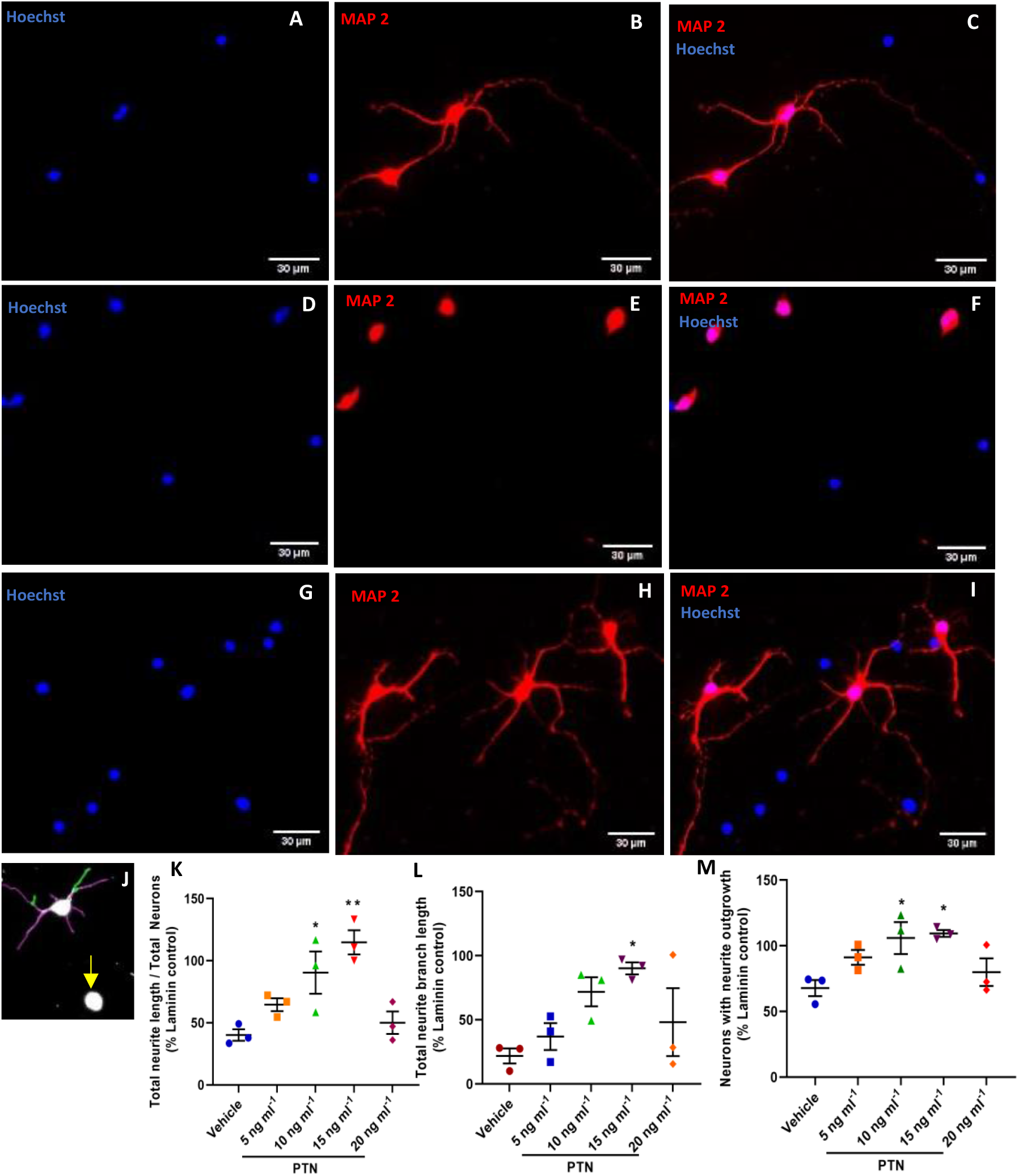
Neurons cultured on laminin (growth permissive matrix) and stained for **(A)** Hoechst, **(B)** MAP 2 (Microtubule associated protein 2), **(C)** overlay. Neurons cultured on laminin + CSPGs (growth inhibitory matrix) and stained for **(D)** Hoechst, **(E)** MAP 2 **(F)** overlay. Neurons cultured laminin + CSPGs (growth inhibitory matrix) and treated with pleiotrophin (PTN) and stained for **(G)** Hoechst **(H)** MAP 2 **(I)** overlay. **(J)** Neurite outgrowth traced using simple neurite tracer plugin (SNT) in image J, total neurite growth (highlighted in purple + green), branched neurite outgrowth (highlighted in green) and neurons with no neurite outgrowth (indicated by yellow arrow). **(K)** PTN induces neurite growth in the presence of CSPGs (ANOVA, *F*_*(4,10)*_=9.028, *P*=0.0024). **(L)** PTN induced branched neurite growth in the presence of CSPGs (ANOVA, *F*_*(4,10)*_=3.740, *P*=0.0413) **(M)** PTN increases number of neurons with neurite growth in the presence of CSPGs (ANOVA, *F*_*(4,10)*_=4.560, *P*=0.0235). **(K – M)** Error bars represent standard error of mean (SEM), symbols *, ** represent p<0.05 and 0.01, respectively, on Dunnett’s multiple comparisons against control. All data are based on 3 independent experiments with a minimum of three technical replicates.

### Pleiotrophin signals via ALK receptor on neurons

PTN signals by inactivating the phosphatase activity of PTPRζ (Herradon and Ezquerra, 2009). Currently available pharmacological inhibitors of PTPRζ are quite limited (Herradõn and Pérez-García, 2014). Thus, identifying other potential receptors for PTN may identify new targets to drive neuron growth. ALK is a putative PTN receptor, and is broadly expressed on cortical neurons *in vitro* (Fig. 2A-C). Treatment of primary neuronal cultures with alectinib, which selectively inhibits phosphorylation of ALK and therefore blocks activity of ALK (Tang *et al*., 2019), was used to probe the involvement of the ALK receptor in PTN signalling. Alectinib induced a dose dependent inhibition of neurite outgrowth in PTN treated neurons cultured on a CSPG matrix (Fig. 2D-J). The available literature indicates the involvement of ALK in neuron like differentiation of PC12 cells (Palmer *et al*., 2009) and depletion of ALK receptor attenuates neuronal proliferation and neurogenesis (Wulf *et al*., 2021). Attenuation of neurite growth due to alectinib treatment provides strong evidence for the necessity of ALK activity in neurite extension, identifying a further avenue for investigation for development of ALK agonists to drive neuroplasticity.

**Figure 2:**
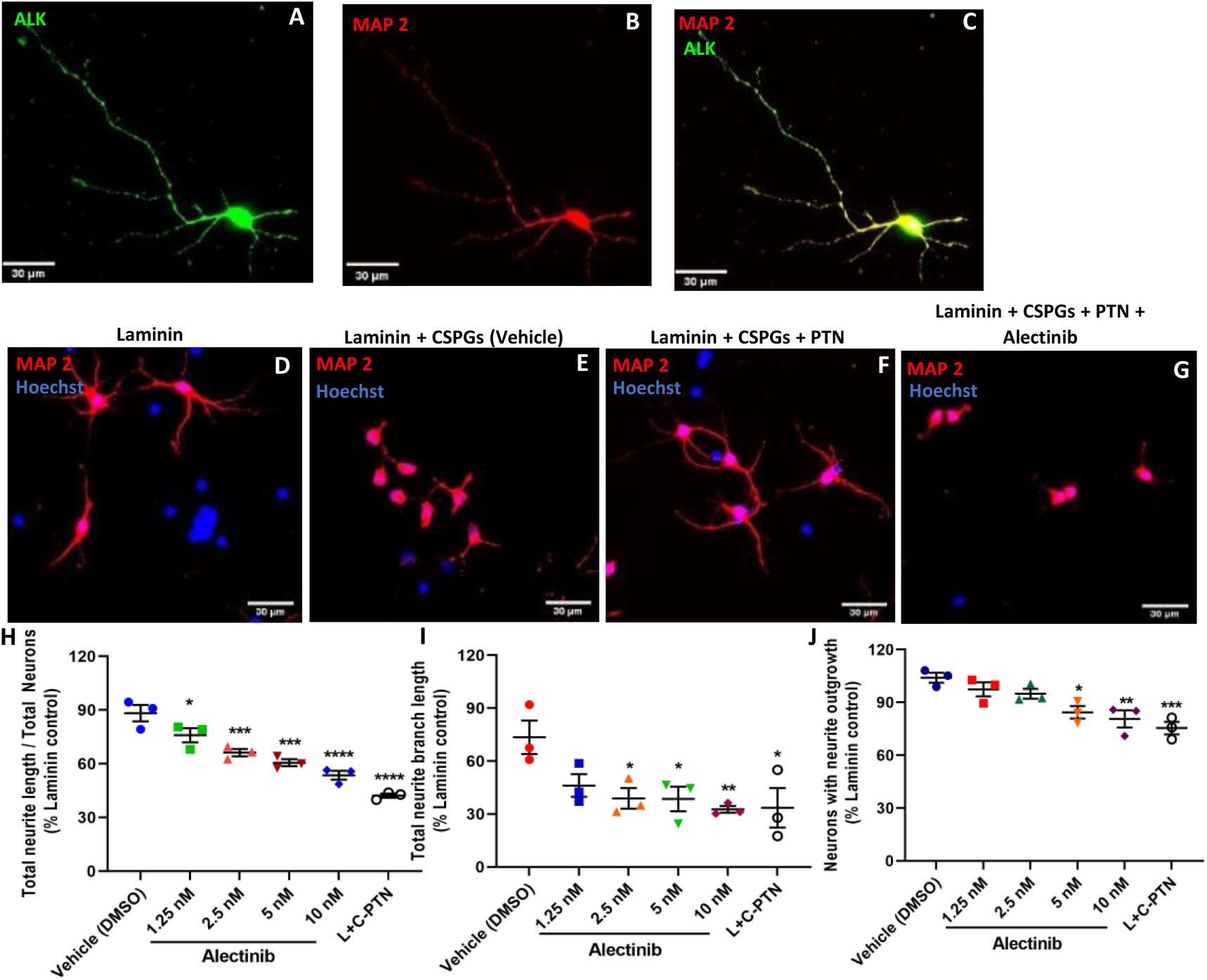
Expression of ALK in cortical neuronal culture and stained for **(A)** ALK **(B)** MAP 2 and **(C)** overlay. Cortical neurons cultured on **(D)** laminin matrix **(E)** Laminin + CSPGs matrix **(F)** Laminin + CSPGs matrix and treated with PTN **(G)** Laminin + CSPGs matrix and treated with PTN and alectinib and stained for MAP2 and Hoechst. **(H)** Neurite growth induced by PTN in the presence of CSPGs is blocked by alectinib (ANOVA, *F*_*(5,12)*_=30.48, *P*<0.0001). **(I)** PTN induced branched neurite growth in the presence of CSPGs is blocked by alectinib (ANOVA, *F*_*(5,12)*_=4.078, *P*=0.0214). **(J)** Alectinib reduced number of neurons with neurite growth in the presence of PTN in the presence of CSPGs (ANOVA, *F*_*(5,12)*_=9.068, *P*=0.0009). **(H – J)** Error bars represent standard error of mean (SEM), symbols *, ** and ***/**** represent P<0.05, 0.01, and 0.001 respectively. All data are based on 3 independent experiments with a minimum of three technical replicates.

### Activating the AKT pathway drives neurite growth

The protein kinase B (AKT) pathway is a key downstream transducer of PTN/ALK signalling (Tang *et al*., 2019). To investigate the involvement of this pathway in the regeneration of neurons, cortical neurons cultured on a CSPG matrix were treated with the AKT activating compound SC79 in the absence of PTN (Tang *et al*., 2019). SC79 treated neurons showed a dose dependent enhanced neurite outgrowth even in the presence of CSPGs (Fig. 3A-G). Although activating AKT signalling with SC79 did not show an elevated response as compared to PTN treatment, with 15 ng mL^-1^ PTN showing 114.8 % and 5uM SC79 with 78.43% increased neurite outgrowth compared to laminin control. The reason for reduced effectiveness of SC79 relative to PTN incubation could reflect other potential downstream activators of ALK signalling. ALK is known to activate many pathways including phospholipase C γ, Janus kinase (JAK), PI3K-AKT, mTOR and MAPK signaling cascades (Herradon and Ezquerra, 2009; Palmer *et al*., 2009). Thus, blocking the ALK activity with alectinib blocks the activity of all these possible pathways and induces a significant reduction in neurite growth even at the lowest tested concentration of 1.25 nM (Figure 2H). PTN/ALK signalling may therefore be a potential target to induce neuron regeneration after CNS injury or degenerative diseases associated with CSPG upregulation, including Alzheimer’s, Parkinson’s, and multiple sclerosis.

**Figure 3:**
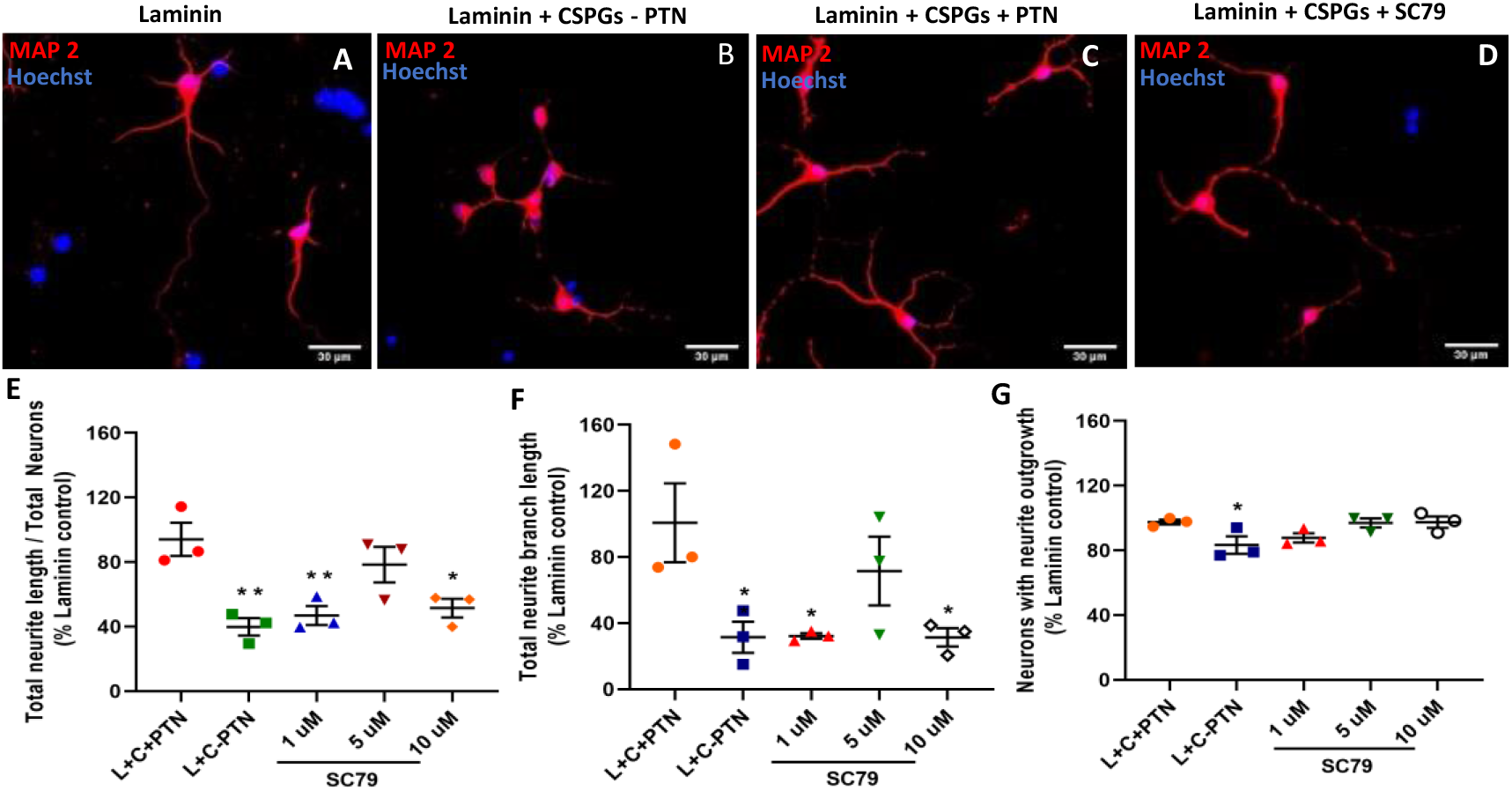
Cortical neurons cultured on **(A)** laminin matrix **(B)** Laminin + CSPGs matrix **(C)** Laminin + CSPGs matrix and treated with PTN **(D)** Laminin + CSPGs matrix and treated with SC79 (AKT activator) and stained for MAP2 and Hoechst. **(E)** SC79 induced neurite growth in the presence of CSPGs (ANOVA, *F*_*(4,10)*_=8.222, *P*=0.0033). **(F)** SC79 increases branched neurite growth of neurons in the presence of CSPGs (ANOVA, *F*_*(4,10)*_=4.441, *P*=0.0254). **(G)** SC79 increases number of neurons with neurite growth in the presence of CSPGs (ANOVA, *F*_*(4,10)*_=3.699, *P*=0.0425). **(E – G)** Error bars represent standard error of mean (SEM), symbols * and ** represent p<0.05 and 0.01, respectively, on Dunnett’s multiple comparisons against vehicle control. All data are based on 3 independent experiments with a minimum of three technical replicates.

## Materials and Methods

### Matrices preparation

Coverslips were coated with 100 µg mL^-1^ Poly-L-Lysine (sigma-aldrich, P5899, USA) for 2 hours. After 3 washes with water, the coverslips were coated with either a growth permissive matrix -10 µg mL^-1^ of laminin (Corning, 354232) or an inhibitory matrix - 10 µg mL^-1^ laminin + 1.25 µg mL^-1^ of CSPGs (sigma aldrich, CC117) for 2 hour and washed 2 times with PBS.

### Primary cortical neuronal culture

All animal protocols were conducted in accordance with Canadian Council on Animal Care Guidelines and approved by the Animal Care and Use Committee: Health Sciences for the University of Alberta. Rat primary cortical neurons were isolated from 0-1day old Sprague Dawley rat pups. The cortices were dissected and digested with TrypLE (Gibco, 12605-028) for 15 minutes at 37°C. The cells were dissociated from tissue by trituration in neurobasal A medium (Thermofisher, 1088802) containing B27 supplement (1:50 v/v) (Gibco, 17504-044), antibiotics and GlutamX (Gibco, 35050-061). The cell suspension was seeded at 20,000 cells / well on coverslips with different matrices and incubated at 37 °C, 5% CO_2_ for 1 hour. Following incubation, the media was incubated (72 h) with media containing different concentration of recombinant human PTN (rhPTN, R&D systems 252 - PL) or the drugs alectinib (Toronto Research Chemicals, C183665) or SC79 (Selleck Chemicals, S7863).

### Immunofluorescence

Cells were fixed after 72 h of treatment with 5% formaldehyde solution for 15 min. and stained with microtubule-associated protein 2 (MAP 2) antibody (1:500, Sigma aldrich M9942), ALK (1:500, abcam ab190934) and Hoechst 33342 (1:1000, Invitrogen, 62249). Statistical analyses of data from epifluorescent images were carried out using one-way ANOVA followed by Dunnett’s multiple comparisons test Using Graphpad Prism (v9.1.2).

## Author contributions

SJG conceived the study with IRW, designed and analysed experiments, and wrote the manuscript. MAC, KGT, and IRW assisted with analysis and interpretation of data and co-wrote the manuscript.

